# BKV clearance time correlates with exhaustion state and T-cell receptor repertoire shape of BKV-specific T-cells in renal transplant patients

**DOI:** 10.1101/556829

**Authors:** Ulrik Stervbo, Mikalai Nienen, Benjamin JD Weist, Leon Kuchenbecker, Patrizia Wehler, Timm H Westhoff, Petra Reinke, Nina Babel

**Affiliations:** Center for Translational Medicine, Medical Clinic I, Marien Hospital Herne, University Hospital of the Ruhr-University Bochum, Herne, Germany; Berlin-Brandenburg Center for Regenerative Therapies, Charité – Universitätsmedizin, Berlin, Germany; Department of Computer Science, Freie Universität, Berlin, Germany; Max Planck Institute for Molecular Genetics, Berlin, Germany

**Keywords:** BKV, T-cell, TCR repertoire, exhaustion, diversity, immunology Abstract

## Abstract

Reactivation of the BK polyomavirus is known to lead to severe complications in kidney transplant patients. The current treatment strategy relies on decreasing the immunosuppression to allow the immune system to clear the virus. Recently, we demonstrated a clear association between the resolution of BKV reactivation and reconstitution of BKV-specific CD4^+^ T-cells. However, which factors determine the duration of viral infection clearance remains so far unclear. Here we apply a combination of in-depth multi-parametric flow cytometry and NGS-based CDR3 beta chain receptor repertoire analysis of BKV-specific T-cells to a cohort of 7 kidney transplant patients during the clinical course of BKV reactivation. This way we followed TCR repertoires at single clone levels and functional activity of BKV-specific T-cells during the resolution of BKV infection. The duration of BKV clearance did not depend on the number of peripheral blood BKV-specific T-cells nor on a few immunodominant BKV-specific T-cell clones. Rather, the T-cell receptor repertoire diversity and exhaustion status of BKV-specific T-cells affected the duration of viral clearance: high clonotype diversity and lack of PD1 and TIM3 exhaustion markers on BKV-specific T-cells was associated with short clearance time. Our data thus demonstrate how the diversity and the exhaustion state of the T-cells can determine the clinical course of BKV infection.

## 1 Introduction

Immunosuppression is one of the most important factors contributing to reactivation of the latent BK polyomavirus (BKV) (1). In fact, BKV reactivation can be observed in up to 80% of all kidney-transplant recipients (2). In as many as 10% of the cases, patients develop BKV-associated nephropathy (BKVAN) which can lead to graft loss (2). Currently, there is no antiviral treatment for BK virus available and the recommended approach to management of BKVAN is a reduction or modification of immunosuppression, in order to achieve sufficient antiviral control by cellular immunity (1,3). In fact, we and others previously demonstrated a strong decrease of the BKV load upon reconstitution of BKV-specific T-cells in renal transplant patients (4,5). Very recently, we demonstrated an improved strategy for monitoring of BKV-specific T-cells by multi-parameter flow cytometry and provided in-depth characteristics of BKV-specific T-cells associated with the initiation of BKV load decline (6).

Although the essential role of BKV-specific T-cells for the initiation of BKV load decline has been well documented, it is so far not clear, which factors determine the clinical course of the disease. The time it takes to clear BKV differs between patients, and the clearance time after severe BKV infection (BKV load more than 100,000 copies/ml) can span weeks to even years (7,8). On the other hand, sustained BKV infection can cause immunopathogenesis and lead to irreversible renal graft tissue damage and graft failure (1). Therefore, understanding the factors that determine the clearance time once BKV load starts declining is crucial for understanding the pathogenesis of BKV-associated nephropathy and for improving therapeutic management in patients with BKV reactivation.

Besides the magnitude and functional activity of antigen-specific T-cells, the heterogeneity of clonal repertoire of antigen-specific T-cells has been suggested to play an important role in the immune defense. Pathogen recognition by T-cells is established through the T-cell receptor (TCR) specific for a cognate epitope in context of the HLA-molecule (9). T-cells with the same progenitor bear identical TCRs and constitute a T-cell clone. Recent studies suggest that diversity and size of the antigen-specific TCR repertoire are two critical determinants for the successful control of chronic infections and the decreased repertoire diversity correlates with the decline of immune responsiveness in ageing (10). Although a young adult human might harbor more than 100 million different TCRs (11), only a small fraction is specific for a given pathogen (12). Despite the low frequency, the identification and tracking of antigen specific TCRs are emerging as valuable tools in the analysis of organ rejection and control of persistent infections (13-15).

In the present work we provide a further characterization of BKV-specific T-cells using the combination of multi-parameter flow cytometry, cell sorting, and NGS-based clonotype profiling. By means of these comprehensive analyses in a small cohort of patients with different course and duration of BKV infection, we identified immunological factors that influence the length of the clearance time. We found a strong correlation between the BKV clearance time and TCR repertoire diversity as well as the expression of exhaustion markers on the surface of BKV-specific T-cells.

## 2 Material and Methods

### 2.1 Study cohort

This study was approved by our local ethical review committee in compliance with the declaration of Helsinki. Informed and written consent was obtained from all patients (Ethic Committee Charité University Medicine, Berlin, Germany, EA2/028/13). The study cohort consisted of 7 kidney transplant recipients with sustained BKV reactivation (Table 1). The HLA typing for each patient and donor is summarized in Figure 1.

**Table 1:** Characteristics of patients with BKV reactivation

| Patient | Gender | Age at Tx<br>(Years) | Donor<br>age | HLA match | BKV Viremia<br>onset after Tx | Immunosuppressive medication |  |
| --- | --- | --- | --- | --- | --- | --- | --- |
|  |  |  |  |  |  | Continual<sup>*</sup><br>(Avg daily dose) | Switch<sup>**</sup><br>(Avg daily dose) |
| 1 | Male | 28 | 54 | 3/6 | 64 days | MMF (1500mg),<br>PRD (4mg) | TAC (13.5mg) →<br>CsA (220mg) |
| 2 | Male | 66 | -- | 2/6 | 122 days | MMF (2000),<br>PRD (4mg) | TAC (7.7mg) →<br>CsA (260mg) |
| 3 | Male | 52 | -- | 3/6 | 32 days | PRD (4mg) | MMF (1080mg) →<br>AZA (75mg);<br>TAC (7.7mg) →<br>CsA (246mg) |
| 4 | Male | 68 | 77 | 4/6 | 94 days | MMF (1500mg),<br>PRD (4mg) | TAC (7.2mg) →<br>CsA (90mg) |
| 5 | Male | 59 | 45 | 0/6 | ~10.24 years | MMF (1750mg),<br>PRD (4mg) | TAC (2mg) →<br>CsA (195mg) |
| 6 | Male | 66 | -- | 0/6 | 67 days | MMF (720mg),<br>PRD (4mg) | TAC (7.5mg) →<br>CsA (195mg) |
| 7 | Male | 65 | 39 | 6/6 | ~6 years | MMF (1000mg),<br>PRD (4mg) | TAC (157mg) →<br>CsA (141mg) |

<sup>\*</sup>Therapeutics that remained constant throughout the course of BKV reactivation. <sup>\*\*</sup> Alteration of therapeutic regiment upon diagnosis of BKV reactivation. Values in parenthesis indicate the average daily dose. Abbreviations: AZA, azathioprine; CsA, Cyclosporine A; MMF, mycophenolate mofetil; PRD, prednisolone; TAC, tacrolimus.

**Figure 1:**
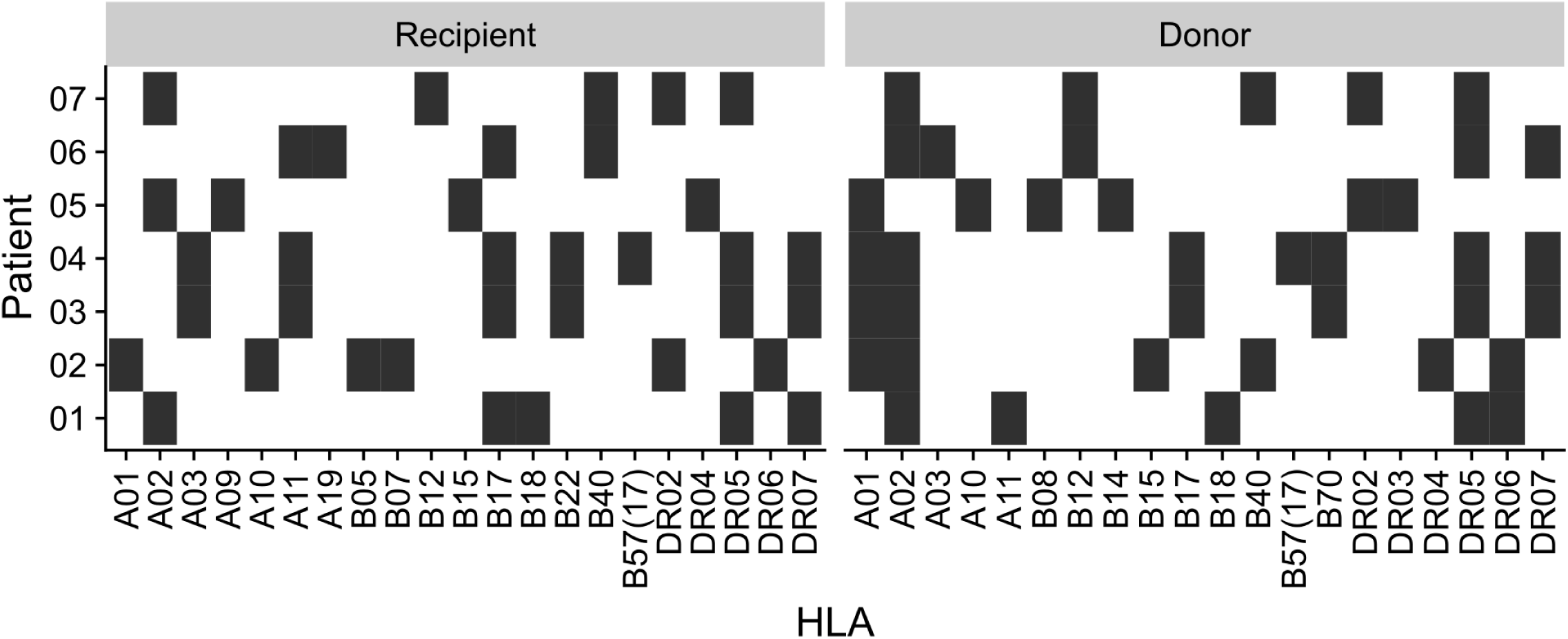
Recipient and donor HLA type. HLA type of the patients and their kidney donors. Black square indicate presence of the HLA type, white indicate absence.

### 2.2 BKV viremia

BKV-DNA copy numbers were determined in serum as previously described (16). Briefly, DNA was isolated from serum using a QIAamp DNA Mini Kit (Qiagen) according to the manufacturer’s instructions. Primers and probes were designed to amplify the VP1 region of BKV. A plasmid standard containing the VP1 coding region was used to determine the copy number per ml. Samples exceeding the detection level (>1000 copies/ml) were considered positive.

### 2.3 BKV stimulation

PBMC were isolated from 40 ml freshly drawn heparin-treated peripheral blood using the Ficoll-Hypaque (PAA Laboratories) gradient method and collected in RPMI-1640 medium (supplemented with L-glutamine (2 mmol/l), fetal calf serum (10%), penicillin (100 IU/ml), and streptomycin (0.1 mg/ml), all from Biochrom) and stimulated with overlapping peptides of all viral products as previously described (17). Briefly, an overlapping peptide pool was created from individual 15mer BKV peptides with an 11 amino acids overlap of the BKV proteins VP1 (Swiss-Prot ID : P14996), VP2 (Swiss-Prot ID : P03094), VP3 (P Swiss-Prot ID : 03094-2), small T antigen (Swiss-Prot ID : P03082) and large T antigen (Swiss-Prot ID : P14999). Although VP3 and small T antigen are included in the ORF of VP2 and large T antigen, respectively, both peptides are included in the overlapping peptide pool to address the shift in the starting position and to mimic the relative abundance of the epitopes. The scheme of the 15mer 11 overlapping peptide pools is illustrated in Figure S1 in Supplementary Material.

All individual peptide pools (JPT) were reconstituted in DMSO and phosphate-buffered saline (PBS) to 1.5 mM (2.5μg/μl) for each peptide. 0.5-1 × 10^7^ PBMCs were incubated with the BKV overlapping peptide pool at final concentration of 1 μg peptide/ml for 16 h at 37 °C, 5% CO_2_. Brefeldin A (Sigma-Aldrich) was added at 1 μg/ml after 2 hours.

### 2.4 Antibodies and staining procedure

Cells were stained with anti-CD3-eF650 (Clone: UCHT1; eBioscience), anti-CD4-PerCP/Cy5.5 (Clone: SK3; BD Bioscience), anti-CD8-BV570 (Clone: 3B5; Invitrogen), and Live/Dead NearIR (Invitrogen). The surface staining was performed in PBS for 15 minutes at room temperature in the dark and subsequently fixed using Permeabilizing Solution 2 (BD Biosciences) for 10 minutes at room temperature.

Intracellular staining was performed with anti-CD137-PE/Cy5 (Clone: 4B4-1; BD Biosciences), anti-CD154-APC/Cy7 (Clone: 24-31; Biolegend), anti-PD1-BV710 (Clone: EH12.2H7; Biolegend), and anti-TIM3-APC (Clone: F38-2E2; eBioscience) for 30 minutes at room temperature in the dark.

Samples were acquired on a LSRII-Fortessa (BD Biosciences) and at least 1.5 × 10^6^ events were recorded. Calibration of the instrument was reconfirmed weekly with rainbow beads (BD Biosciences).

### 2.5 TCR repertoire analysis

T-cells were sorted by flow cytometry according to CD4 and CD8 expression from unstimulated PBMCs stained with anti-CD3-eF650 (Clone: UCHT1; eBioscience), anti-CD4-PerCP/Cy5.5 (Clone: SK3; BD Bioscience), anti-CD8-BV570 (Clone: 3B5; Invitrogen), and Live/Dead NearIR (Invitrogen) at the BCRT Flow Cytometry Laboratory. For isolation of BKV-specific T-cells, PBMCs were stimulated with the BKV overlapping peptide pool at final concentration of 1 μg peptide/ml for 16 h at 37 °C, 5% CO_2_.in the presence of 1μg/ml CD40 (Miltenyi Biotec). Activated BKV-specific T-cells were MACS isolated using the IFNγ Secretion Assay – Cell Enrichment and Detection Kit, human (Miltenyi Biotec), per manufacturer’s instructions. Genomic DNA was isolated using the AllPrep DNA/RNA Micro Kit (Qiagen).

The recombined V-CDR3-J region of the recombined genomic TCR-β locus was amplified with partly degenerate primers covering all functional Vβ and Jβ regions (0.25 μM each) as previously described (18). Using Phusion Hot Start II DNA Polymerase (Thermo Fisher) and DNA template (up to 1 μg) in a final volume of up to 100 μl, the PCR reaction was heated to 98°C for 5 minutes and followed by typically 20 cycles of 98°C 15s, 65°C 30s and 72°C 30s, with final extension at 72°C for 5 min. The PCR product was purified with Sera-Mag SpeedBead Carboxylate-Modified Magnetic Particles (GE Healthcare Life Sciences) and after wash with 70% EtOh resuspended in nuclease-free water (Thermo Fisher). Purified PCR product was used in a second PCR where sample indices were added as above, with only 14 cycles of PCR. Indexed PCR product was separated on 2%-LMP agarose (Sigma) and purified with the Gel Extraction Kit (Qiagen).

Sequencing library preparation with consequent sequencing was performed using Illumina MiSeq Technology at the Next Generation Sequencing Core Unit at Berlin Brandenburg Center for Regenerative Therapies, Berlin, Germany. Reads with an average quality score below 30 were excluded from further analysis. The remaining high quality reads were processed using IMSEQ to identify the CDR3 amino acid sequence (19). CDR3 embedded at wrong reading frames and with stop codons were discarded. On average we obtained 488,700 reads per sample, with first and third quantile of 279,400 and 626,400, respectively. Antigen specific clonoypes of the unstimulated whole-blood samples were identified by overlap to clonotypes obtained by the IFNγ Secretion Assay.

### 2.6 Diversity estimation

To account for differences in sequence depth, the clonotypes were rarified to equal sample size before estimation of repertoire diversity. Rarefaction was performed using the R-package vegan, version 2.4-3 (20). The Shannon index (21) is given by

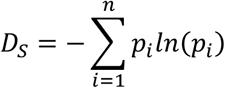

where *p*_*i*_ is the proportion of the *i*th clonotype in the population.

The Berger-Parker index (22) is given by

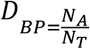

where N_A_ indicates the abundance of the most abundant clonotype, N_T_ is the total number of clonotypes. This index relies on the most abundant clonotype and is therefore less sensitive to sequencing errors.

### 2.7 Estimation of clearance time

Patients were considered free from BKV viremia if at least two consecutive measurements were below detection limit. The time from the highest viral load until complete virus clearance was estimated from the slope of the linear fit of the log-transformed measurements spanning from the highest BKV copy number to the first BKV viremia free measurement. The initial conditions for the clearance time estimation were the highest measured viremia level:

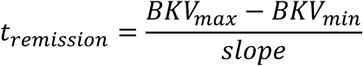

where BKV_min_ is the detection limit of 1000 BKV copies/ml.

### 2.8 Statistical analysis

Flow cytometry data was analyzed using FlowJo version 10.3 (Tree Star). Statistical analysis was performed using R, version 3.3.1 (23). p values < 0.05 were considered significant.

## 3 Results

### 3.1 BKV viremia clearance time differs between kidney transplant recipients

BKV viremia was monitored in kidney transplant patients with BKV reactivation (Figure 2). The highest measured BKV copy number followed by load decline was taken as the start of the viremia clearance phase. The patient was considered free from BKV viremia when samples from at least two consecutive time points were both negative. The first of these time points marked the end of the viremia clearance phase. The actual clearance is at a point before the observed clearance. To limit the effect of sampling frequency, we sought to obtain a data driven length of the viremia clearance phase by linear regression on log-transformed BKV viral load. The slope of the fitted curve was used the slope to estimate the number of days until the virus was cleared. The linear fit explained 78 to 96% of the variation in the viremia decline phase (Figure 2A). The estimated clearance time ranged between 46 and 298 days (Figure 2B). The time with increasing BKV viremia was similarily estimated through linear regression (Figure 2C). No association bethween the length of the inclining and declining phase was found (data not shown).

**Figure 2:**
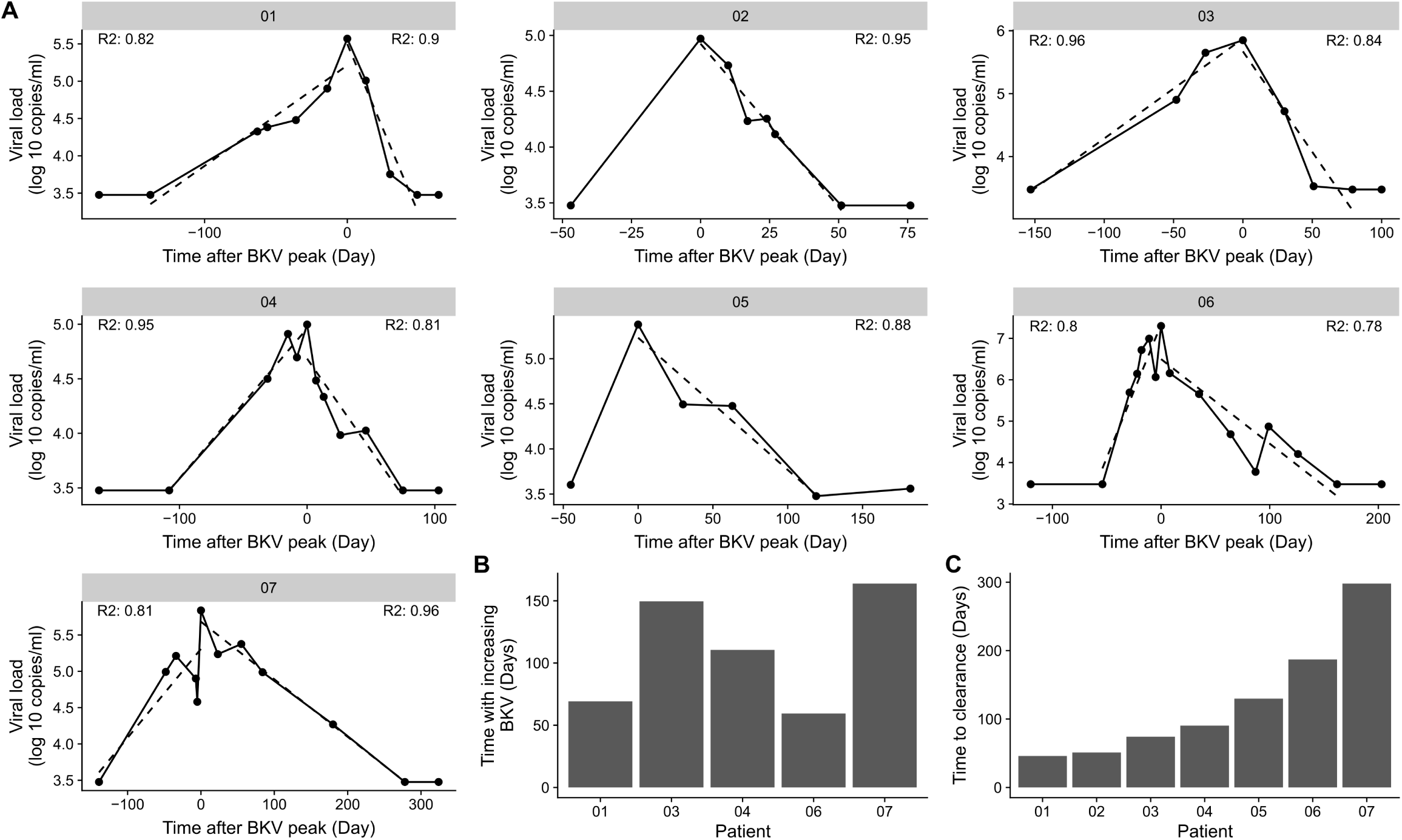
Difference in BKV viremia clearance time. A) BKV viremia remission phase for the individual patients in the cohort. The dotted line indicates the best linear fit from the time point before BKV-reactivation to highest BKV load and from the highest BKV load to the first time point with BKV viremia below detection limit. R^2^ indicate the goodness of fit for the time of increasing BKV viremia and for the time with decreasing viremia. B) Time with increasing BKV viremia for each patient arranged by the time to clear the virus. C) Clearance time calculated from the slope of a straight line from the highest BKV value to the first time point with BKV viremia below detection limit.

### 3.2 Magnitude and phenotypic characteristics of BKV-specific T-cells do not explain differences in BKV clearance time

Previously, we showed that the BK viral load decline was associated with the reconstitution of helper and cytolytic BKV-specific T-cells in peripheral blood (6). However, the factors that affect the BKV infection clearance time were so far not addressed. Here, we analyzed the effect of functional and phenotype characteristics on the duration of BKV clearance (Figure 3A, Figure S2-S8 in Supplementary Material). First, using the previously published gating strategy and activation markers CD137/CD154 for CD4^+^ T-cells and CD137/Granzyme B for CD8^+^ T-cells (6), the frequencies of BKV-specific CD4^+^ and CD8^+^ T-cells circulating in peripheral blood were evaluated. No correlation between the duration of BKV clearance time and the ratio of either BKV-specific CD4^+^ or CD8^+^ T-cells at the peak of viral load could be observed (Figure 3B and C). The frequencies of total CD4+ or CD8+ also did not correlate with the clearance time (Figure S9A and B in Supplementary Material).

**Figure 3:**
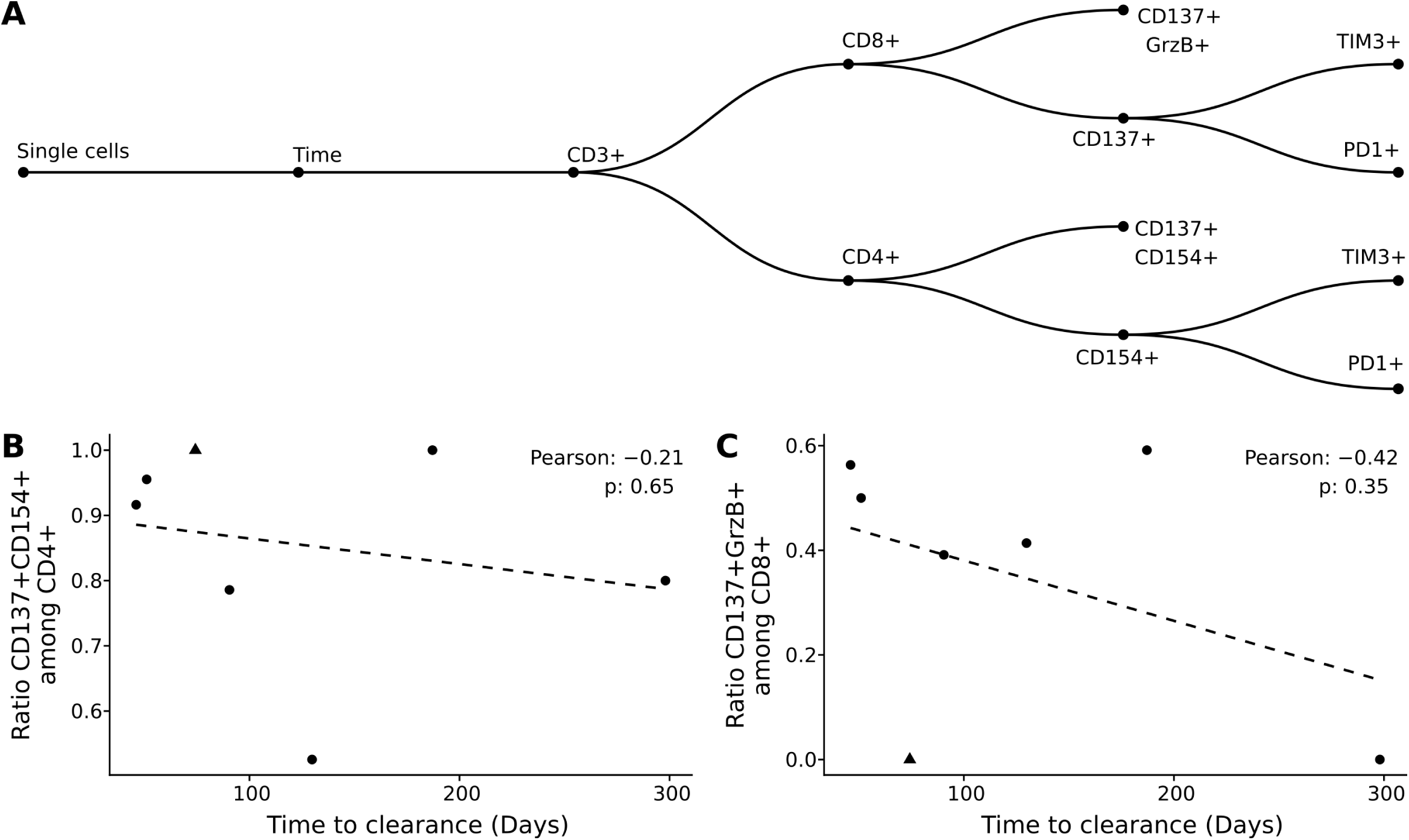
Clearance time is not explained by magnitude and phenotypic characteristics of BKV-specific T-cells. A) Gating strategy applied in the current study. B-C) Pearson correlation of clearance time with BKV-specific CD4^+^ T-cells (B) and BKV-specific CD8^+^ T-cells (C). Each point represents a patient in the study and the dotted line the best linear fit.

Since previous data demonstrated a better antiviral control in patients with elevated number of multi-functional cells (17), we next addressed the question whether functionality of BKV-specific T-cells could explain the difference in clearance time. To this end we analyzed multi-functional characteristics of BKV-specific T-cells as defined by multiple cytokine production (IL2, TNFα and IFNγ) and/or effector molecule Granzyme B. The analysis of BKV-specific cytokine multifunctional and effector molecule producing capacities did not reveal any influence on duration of clearance period (Figure S9C and D in Supplementary Material).

### 3.3 Tracking of BKV-specific TCRs

Previous studies on the protective function of cellular immunity in cancer suggest an important role of the TCR repertoire (24,25). However, in case of viral infections it is not clear, whether few immunodominant clones or rather diverse, polyclonal TCR repertoires determine the clinical course infection (26,27). Addressing this question, we analyzed BKV-specific T-cells on a single clone level during BKV clearance.

We have previously used NGS-based clonotype profiling to identify BKV-specific T-cell clones and track them in peripheral blood, renal tissue biopsy, and urine (18). Using a similar approach, we were able to study the kinetics of BKV-specific T-cells on the clonal level of whole blood-derived CD4^+^ and CD8^+^ T-cell subsets in the current study (Figure 4). Antigen specific clonoypes within whole-blood samples were identified by identity to BKV-specific clonotype sequences obtained by the IFNγ Secretion Assay. The distribution of length TCRβ CDR3 of these BKV-specific clonotypes had a similar shape for all patients and was normally distributed in all cases (Figure S10 in Supplementary Material). A tendency for slightly longer CDR3 was observed for patient 3 (17 AA on average compared to 15 AA for the others; Table S1 in Supplementary Material). However, the most abundant length for the patients was 15 AA.

**Figure 4:**
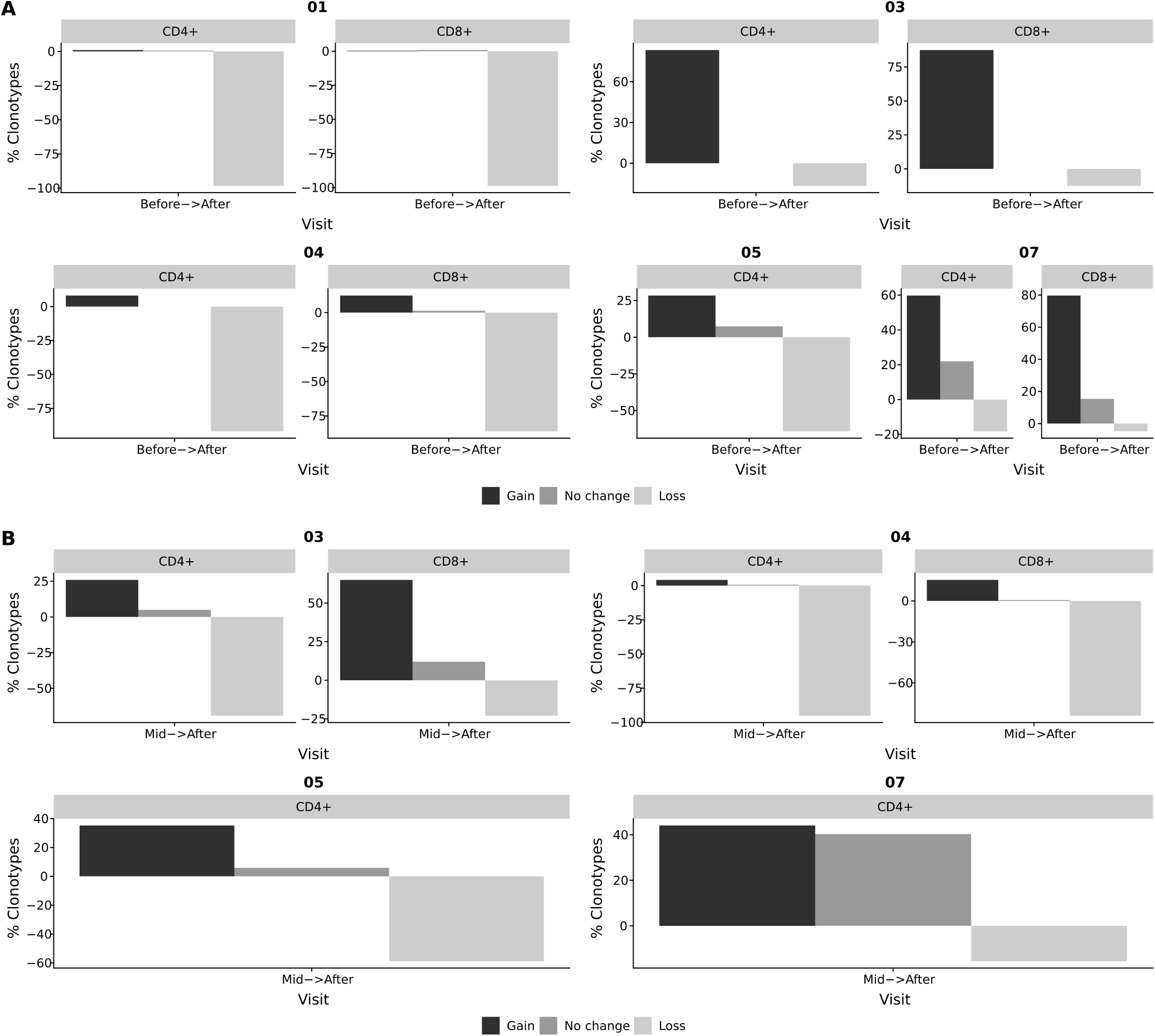
Tracking of BKV-specific TCR clonotypes. BKV-specific TCR clonotypes were obtained from IFNγ producing T-cells after stimulation with BKV overlapping peptide pools. TCR clonotypes from whole blood CD4^+^ and CD8^+^ subsets were obtained at different stages of viral clearance. The overlap between the clonotypes of whole blood samples and clonotypes of the IFNγ producing T-cells were identified as BKV-specific T-cells in circulation. The abundance of these circulating BKV-specific T-cells were subsequently compared at distinct stages of viral clearance. Black bars indicate the frequency of clonotypes gained from the earlier to the later time point. Dark grey indicates the frequency of clonotypes found at both time points. Light grey indicates the frequency of clonotypes that has disappeared from the earlier time point to the later. A) Relative change in BKV-specific clonotypes from the initiation of viral clearance (before) to the resolution of BKV infection (after). B) Relative change in BKV-specific clonotypes from T-cells obtained during the clearance phase (mid) to the resolution of BKV infection (after). The bars indicate the frequency of clonotypes gained, lost, or sustained as the viral clearance progressed. Gained clonotypes – clonotypes that were present after viral clearance, but not before; lost clonotypes – clonotypes that were present at the beginning of the clearance phase but not thereafter; and sustained clonotypes – clonotypes present before and after viral clearance.

The TCRβ V-segment V6 dominated the BKV-specific clonotypes in all patients. The segment V7 was also dominant in the patients 1 and 7 (Figure S11A in Supplementary Material). The J-segment J2-1 was found to be frequent in all patients, but also J1-1 and for patient 5 the J1-6 were also present (Figure S11B in Supplementary Material). When comparing the V-J-segment usage, combinations of the most dominant V- and J-segments for each patient were frequently found, but otherwise no particular pattern was visible (Figure S11C in Supplementary Material).

In our study we identified BKV-specific T-cell clonotypes and followed them in peripheral blood of renal transplant patients at the time point of initiation of viral decline and after the resolution of BKV infection. While tracking BKV-specific clonotypes during the course of viral decline we observed substantial changes in TCR repertoires. While only 0.9% (range: 0 – 22.0%) of clonotypes remained stable during the clinical course of BKV clearance, 28.4% (range: 0.4 – 87.5%) of clonotypes showed expansion and 64.2% (range: 4.9 – 98.8%) of clonotypes showed loss in their frequencies (Figure 4A). We also analyzed changes in TCR repertoire between mid-time of viral decline and after BKV clearance (Figure 4B). Similarly strong repertoire changes were observed in most patients in the course of viral load decline (mid-time – resolution). With respect to the role of repertoire turn-over for the viral clearance, no specific patterns could be clearly detected. However, in patient with severe long-lasting BKV replication (patient 7), we observed a high number of stably detectable clonotypes (40 % of BKV-specific clones). This case was especially pronounced between mid-time and viral clearance as compared to other patients (Figure 4B).

### 3.4 Low diversity of BKV-specific clonotypes indicates prolonged clearance time

We next asked how the TCR repertoire diversity of BKV-specific T-cells reflected the differences in clearance time. We first evaluated the commonly used Shannon index, but found no pattern associating with the clearance time (Figure 5A). Berger-Parker is another diversity index, which disregards the clonotypes with very low frequency (that is the tail of the distribution), and can therefore be considered a more robust index (28). When we looked at inverse of the Berger-Parker index – which directly reflects the clonal diversity of an analyzed population – we found a strong correlation to the clearance time (Figure 5B, Figure S12D in Supplementary Material; Pearson correlation: −0.99, p-value: 0.013).

**Figure 5:**
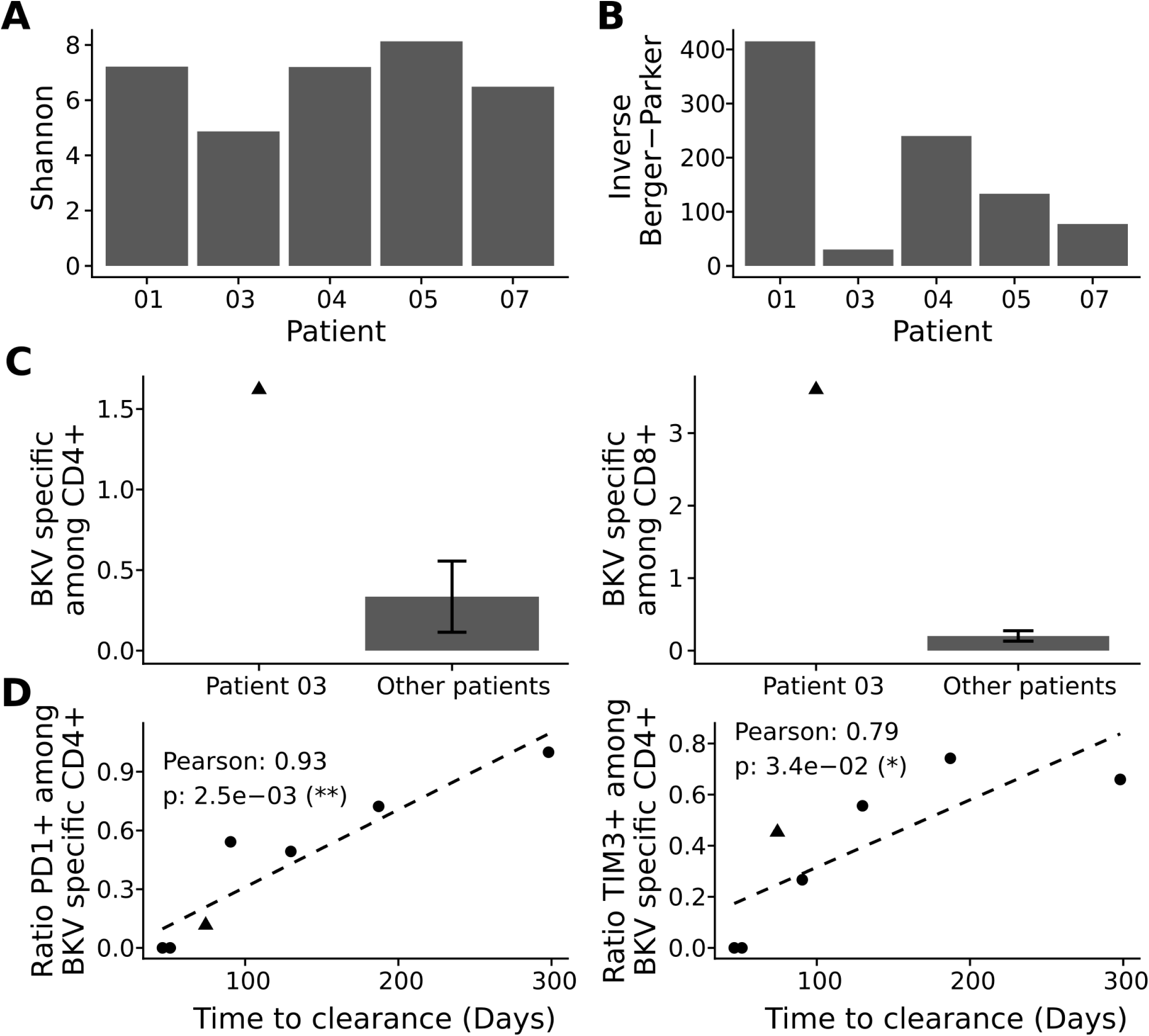
Repertoire analysis of BKV-specific T-cells. BKV-specific T-cells were isolated and subjected to clonotype analysis. A) Population diversity by the Shannon index (A) and the inverse Berger-Parker index (B). C) Comparison of total BKV-specific CD4^+^ T-cells (left panel) and CD8^+^ T-cells (right panel), for patient 3 and the other patients. D) Pearson correlation of clearance time and cellular exhaustion as marked by PD1 and TIM3 among activated CD4^+^ T-cells, see Figure S2-S8 in Supplementary Material for details. Presented is the ratio of stimulated cells to negative control (DMSO treated). Each point represents a patient in the study and the dotted line the best linear fit.

We also assessed the clonal size of the TCR repertoire and cumulative frequencies of BKV-specific TCR by analyses of most abundant BKV-specific T-cell clones (Figure S12E in Supplementary Material). Clonal size data were in line with the data on repertoire diversity. Thus, in cases of low diversity, cumulative frequencies of the most abundant clonotypes were much higher and accounted for a lower diversity than in cases where cumulative frequencies of most abundant clonotypes were low. In detail, the lowest repertoire diversity corresponded to 25.1 % as a cumulative frequency of 10 most abundant clones, whereas the highest repertoire diversity corresponded to a cumulative frequency of 2.0% for the 10 most abundant clones. Accordingly to the diversity index analyses, low cumulative levels correlated with short clearance time (Pearson: 0.99, p-value: 0.01). The observed correlations were not due to general differences in complete CD4^+^ T-cell repertoire (Figure S12A-C in Supplementary Material).

### 3.5 Activation can counteract low diversity

Patient 3 clearly rejected the observed pattern. This particular patient showed a fast BKV load decline but a very low level of BKV-specific TCR repertoire diversity (Figure 5B). We therefore asked whether the expansion of few BKV-specific clones could explain this aberrant behavior. We reasoned that any expansion *in vivo* would be reflected in expanded levels of antigen specific T-cells and performed, consequently, in-depth characterization of BKV-specific T-cell immunity in this patient by flow cytometry. Of interest, we detected very high frequencies of BKV-specific CD4^+^ and CD8^+^ T-cells in peripheral blood of patient 3; the magnitude of the T-cell response was almost 10 times higher compared to the frequencies among the other patients (Figure 5C).

### 3.6 Prolonged clearance time correlates with expression of exhaustion markers

In addition to the demonstrated correlation between repertoire diversity and clearance time, we also evaluated the role of functional exhaustion of BKV-specific T-cells. Flow cytometric analysis of the exhaustion markers PD1 and TIM3 on CD4^+^ T-cells demonstrated a very strong correlation between the expression of these markers on BKV-specific CD4^+^ T-cells and the clearance time (Figure 5D). Interestingly no association to co-expression of PD1 and TIM3 was observed (Figure S13C in Supplementary Material). No expression of either exhaustion marker was seen among BKV-specific CD8^+^ T-cells (Figure S13A and B in Supplementary Material). Our data thus show that exhaustion of the CD4^+^ T-cells is associated with extended clearance time.

## 4 Discussion

The present study provides characterization of BKV-specific T-cells in a small cohort of renal transplant recipients. In line with previous reports (7,8), we observed significant differences in clearance time of BKV virus. We used modern high throughput technologies such as next generation sequencing and multi-parameter flow cytometric analysis to address the question how the fitness of the immune system affects the BKV clearance time. Here, we presented evidence that the diversity of the BKV-specific TCR repertoire inversely correlates with the duration of BKV clearance from initiation of the load decline until virus clearance. In addition, we showed that the expression of exhaustion markers PD1 and TIM3 on BKV-specific CD4^+^ T-cells at the time point of initial BKV load decline correlates with sustained BKV reactivation, while lack of the PD1 and TIM3 expression corresponded to shorter clearance time. These data therefore suggest that the repertoire diversity and functional fitness of BKV-specific T-cells as defined by the expression of exhaustion markers are key players in determining the clinical course of viral infection and clearance.

The essential role of T-cells in controlling BKV reactivation and clearance is well established. We have previously shown that the loss of BKV-specific T-cells is associated with a higher risk of BK viremia (29) and that an increase in BKV-specific T-cell response corresponded to the clearance of BKV reactivation (4). The current recommendation for the management of BKV infection in renal transplant patients includes accordingly a reduction or modification of immunosuppressive therapy allowing for an immune system reconstitution. In fact, frequencies of BKV-specific T-cells increase upon reduction of immunosuppression as demonstrated previously (5,30). However, despite similar therapeutic approaches, the duration of BKV clearance varies among patients spanning period of time from several weeks to even years (7,8). Factors that might be responsible for the different clinical course of BKV infection are not identified so far. Addressing the role of the magnitude of BKV-specific T-cell immunity we analyzed the frequencies of BKV-specific T-cells in a small patient cohort. No correlation was found between the duration of viral clearance and the frequencies of BKV-specific T-cells. We also analyzed the role of the functional characteristics of BKV-specific T-cells on the clinical course of infection. Previous studies demonstrated an important role of multi-functional T-cells for the control of chronic infections (31-33). Our own data in patients with a history of resolved low-level BKV infection demonstrated a higher number of multi-functional CD4^+^ T-cells compared to patients with a history of severe long-lasting BKV infection (17). Our present finding that the frequencies of BKV-specific multi-functional T-cells at the peak of viral load did not associate with a shorter clearance time, indicate the importance of other factors once the viral control is broken.

Assessment of T-cell receptor repertoires and T-cell clonality is the focus of many immunological studies providing in-depth T-cell characterization (24,34-36). This technology provides a unique potential for the analysis of memory T-cells in humans, e.g. clonal relations between circulating and resident cells or various functional subsets (34,37); assessment of T-cell repertoire size and diversity(10); detection and tracking of antigen-specific T-cells within various tissue samples (14,18). In some clinical situations, TCR repertoire profiling is the only available method to differentiate between an undesired pathological T-cell response and a local protective antiviral immune response, i.e. the case of acute transplant rejection by alloreactive T-cells versus BKV clearance by virus-specific T-cells infiltrating the kidney transplant (18,38). Questions regarding the role of clonotype diversity and protective capacity of the adaptive cellular immune response remain so far not completely resolved. Some studies suggest that diversity and size of the antigen-specific TCR repertoire are two critical determinants for successful control of chronic infections and that the decreased repertoire diversity in age correlates with the decline of immune responsiveness in ageing (10).

Although it is generally believed that a large T-cell clonotype diversity provides the optimal protection (27,39), mice with reduced TCR diversity are capable of controlling LCMV or Sendai virus infections (40) and differences in diversity of influenza-specific CD8^+^ T-cells were not reflected in differences in overall functionality (41). On the other hand, an association between TCRβ chain high diversity and low incidence of CMV and EBV reactivations has been observed in solid organ or hematopoietic stem cell transplant patients such that higher diversity yields lower incidence (42-44).

In line with these data, we here demonstrated a negative correlation between diversity level and clearance time of BKV infection. However, a single patient defied the observed association. We believe that the low diversity level was substituted by a high level of functional fitness of BKV-specific T-cells. In fact, we observed a nearly 10 times higher frequency of BKV-specific T-cells in the analyzed as compared to the other patients and very low levels of PD1 or TIM3 expression on BKV-specific CD4^+^ T-cells at the peak of BKV viral load. The large frequency and low diversity was reflected in a strong dominance of the first ten most abundant BKV-specific clonotypes which indicates an expansion of particular T-cell clones. The relationship between the strength of the TCR signal and the proliferation of the T-cell is well established (45). It is therefore possible that highly specific T-cell clones dominate in this one patient, resulting in low diversity of the overall BKV-specific repertoire. In a more general sense, it might be that most BKV-specific TCRs are of lower avidity and that high avidity TCRs are rare and generated by chance. This also explains the observation, where low diversity leads to longer clearance times: a broader TCR repertoire is better equipped to generate a population of BKV-specific T-cells required to initiate and maintain the clearance of the virus (4,5). We are currently in the process of elucidating the avidity of a select number of the identified BKV specific clonotypes.

An alternative explanation to the low diversity but fast clearance of patient 3 could be that the patient has a particular set of HLA molecules, which allow strong presentation of immunogenic BKV epitopes. However, the haplotypes of patient 3 were the same as for patient 4; at least on the level of allele group. This indicates that the HLA haplotype alone is insufficient to explain the TCR diversity.

The expression of the exhaustion markers PD1 and TIM3 on the CD4^+^ T-cells displayed in both cases a straight forward relationship with the clearance time, such that the absence of exhaustion markers was associated with shorter clearance time. This makes it possible, that the fitness of the T-cells is a major player in creating a pool of BKV controlling T-cells in addition to the repertoire diversity. Since both the repertoire diversity and the expression of exhaustion markers correlated with the clearance rate, it is possible that one effect depends on another one: low diversity leads to insufficient clearance and therefore to long-lasting BKV reactivation with a subsequent sustained activation of BKV-specific T-cells. This long-lasting T-cell activation could result in a high-level of exhaustion. Alternatively, the expression of exhaustion makers leading to a functional impairment of T-cells might lead to a protracted viral elimination and contribute therefore to a permanent activation of BKV. Such permanent antigenic stimuli can lead to clonal skewing with an expansion of a few clones, similar to a phenomenon described for older patients with CMV reactivations (46). Although our study provided first observations between T-cell functionality, duration of viral clearance, and T-cell clonality (Figure 6); the interplay between the T-cell exhaustion status and the TCR repertoire diversity is of special interest and will be addressed in ongoing studies.

**Figure 6:**
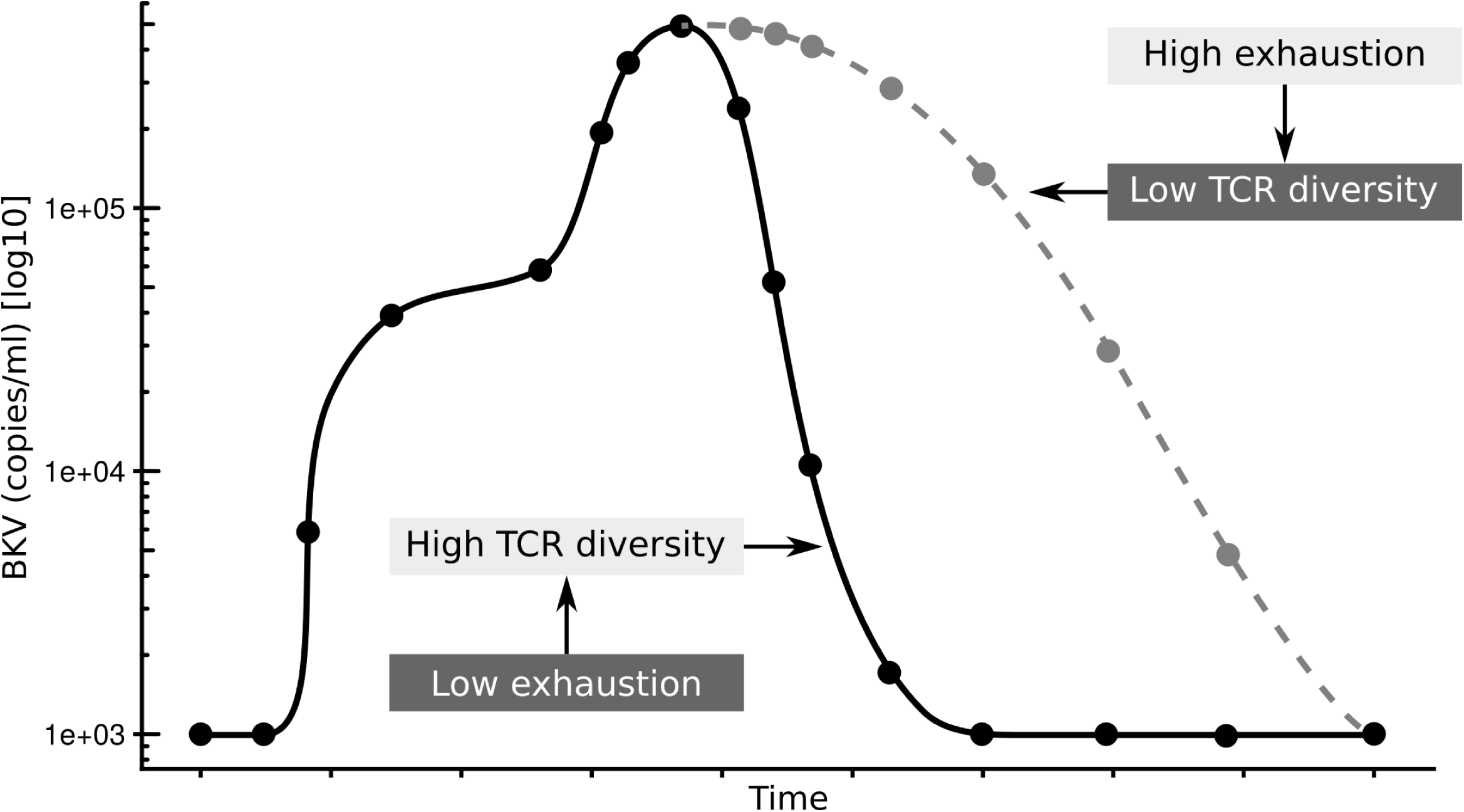
Summary of hypothesis. Low exhaustion and high TCRβ diversity results in shorter clearance time (black line), while high exhaustion and low TCRβ diversity results in shorter clearance time (gray line).

Our study is limited in that it is based on a small cohort of 7 kidney transplant patients, with TCRβ CDR3 data available for only 5 of these 7 patients. This is arguably a very small cohort. However, the observed associations had high correlation coefficients indicating small variance in the data, and thus demonstrating that the findings presented here are robust, despite the small cohort. The associated p-values were likewise small, confirming that the observations might not be per chance only. The viral load can fluctuate by up to one log, but we generally observed a steady decline in BKV copies/ml after some maximal value. We therefore defined the start of the clearance phase as the peak in BKV viremia. Other definitions might be valid, but they would affect the clearance duration for all patients equally, and thereby not disturb the order of clearance times.

Though the results presented in our study are very encouraging, the small cohort size warrants caution. The second limitation is the sampling schedule. Since the course of the viral reactivation is inherently unpredictable, the acquisition of samples at the very early stage/and or peak of reactivation is very much a case of “hit and miss”. Future studies on a large patient cohort addressing the role of TCR diversity and T cell exhaustion in general are required to validate our findings. To include the most relevant periods of BKV reactivation and make such monitoring schedule logistically feasible, blood should be collected and stored as isolated and frozen PBMCs at regular intervals. In summary, we propose that the exhaustion status of CD4^+^ T-cells affects the diversity of the antigen specific TCR repertoire resulting in prolonged clearance time after BKV reactivation (Figure 6). It will be interesting to see if this is a particular feature of the control of BKV in a transplantation setting, or if this is general to chronic infections.

## 6 Author Contributions

US, analyzed the data, created the figures and drafted the manuscript; MN performed the experiments and analyzed the data; BW and WP performed the experiments; LK analyzed the data; TW, PR, and NB designed the study; all authors revised the manuscript and approved its final version.

## 8 Acknowledgments

We would like to acknowledge the excellent assistance of the BCRT Flow Cytometry Laboratory and of the BCRT Next Generation Sequencing Core Unit. We acknowledge support from the Federal Ministry of Education and Research (BMBF) and the Open Access Publication Fund of Charité – Universitätsmedizin Berlin.

## 9 Data Availability Statement

The raw data supporting the conclusions of this manuscript will be made available by the authors, without undue reservation, to any qualified researcher.

