## Supplemental Information for "BKV clearance time correlates with exhaustion state and T-cell receptor repertoire shape of BKV-specific T-cells in renal transplant patients"

### 1 Supplementary Figures and Tables

#### 1.1 Supplementary Figures

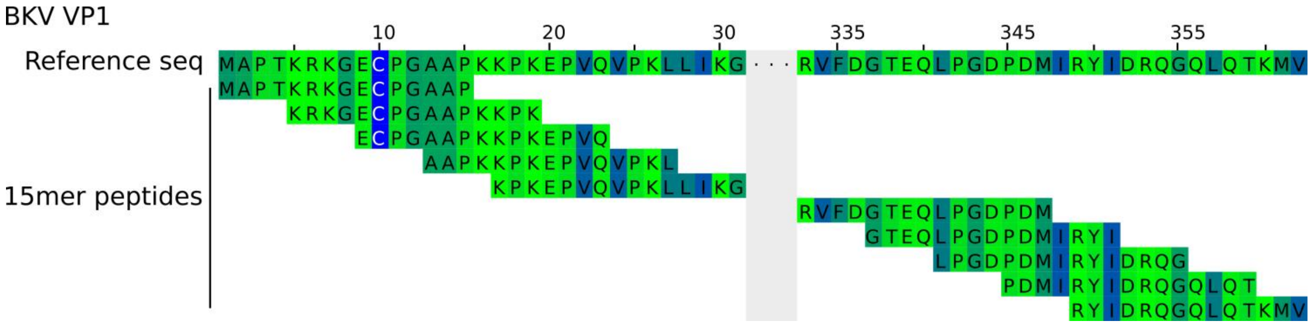


**Supplementary Figure 1.** Principle of 15mer 11 AA overlapping peptides from BKV VP1 peptide sequence. The entire BKV VP1 peptide is 362 AA long, but is presented here with position 32 to 332 truncated (gray area).

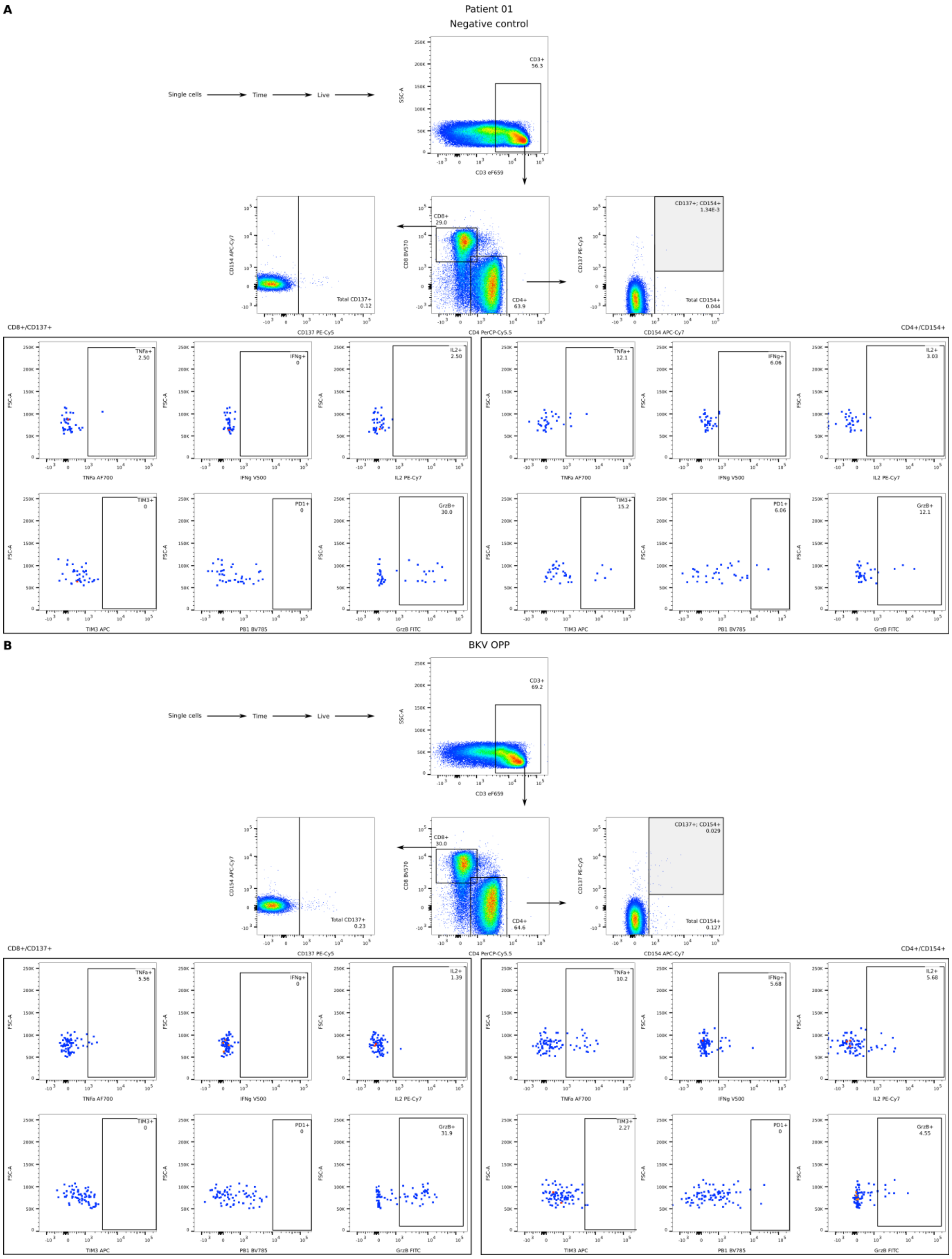


**Supplementary Figure 2.** Gating strategy for the phenotypic and functional characterization of BKV-specific T cells. Complete gating strategy from identification of CD3$^+$ T cells to the assessment of cytokines and exhaustion markers is presented for patient 01. A) Negative DMSO control. B) Stimulation with BKV OPP. For CD4$^+$ CD154$^+$CD137$^+$ (gray box) and total CD154$^+$ (gray and white box) were identified.

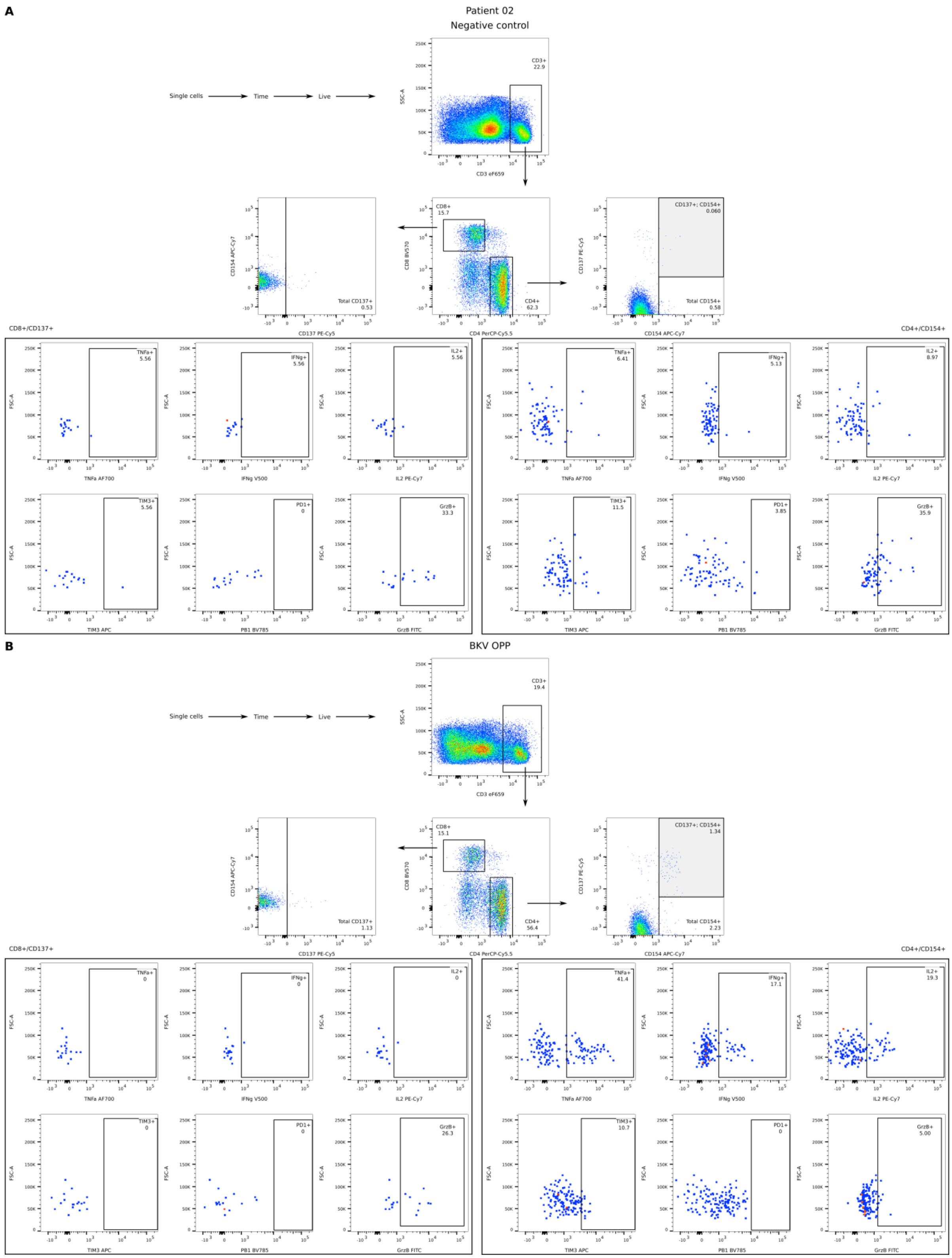


**Supplementary Figure 3.** Gating strategy for the phenotypic and functional characterization of BKV-specific T cells. Complete gating strategy from identification of $CD3^{+}$ T cells to the assessment of cytokines and exhaustion markers is presented for patient 02. A) Negative DMSO control. B) Stimulation with BKV OPP. For $CD4^{+}$ $CD154^{+}CD137^{+}$ (gray box) and total $CD154^{+}$ (gray and white box) were identified.

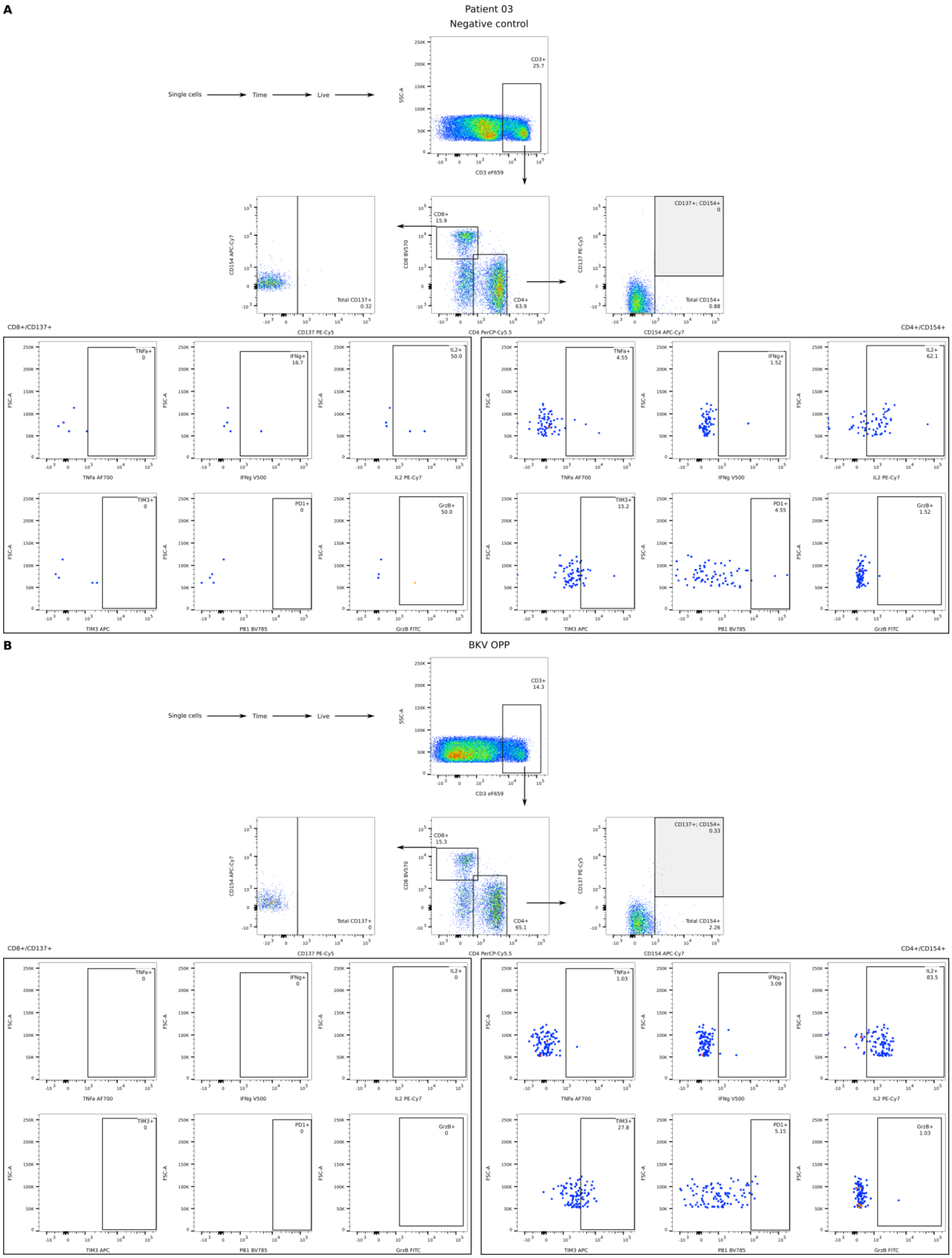


**Supplementary Figure 4.** Gating strategy for the phenotypic and functional characterization of BKV-specific T cells. Complete gating strategy from identification of $CD3^{+}$ T cells to the assessment of cytokines and exhaustion markers is presented for patient 03. A) Negative DMSO control. B) Stimulation with BKV OPP. For $CD4^{+}$ $CD154^{+}CD137^{+}$ (gray box) and total $CD154^{+}$ (gray and white box) were identified.

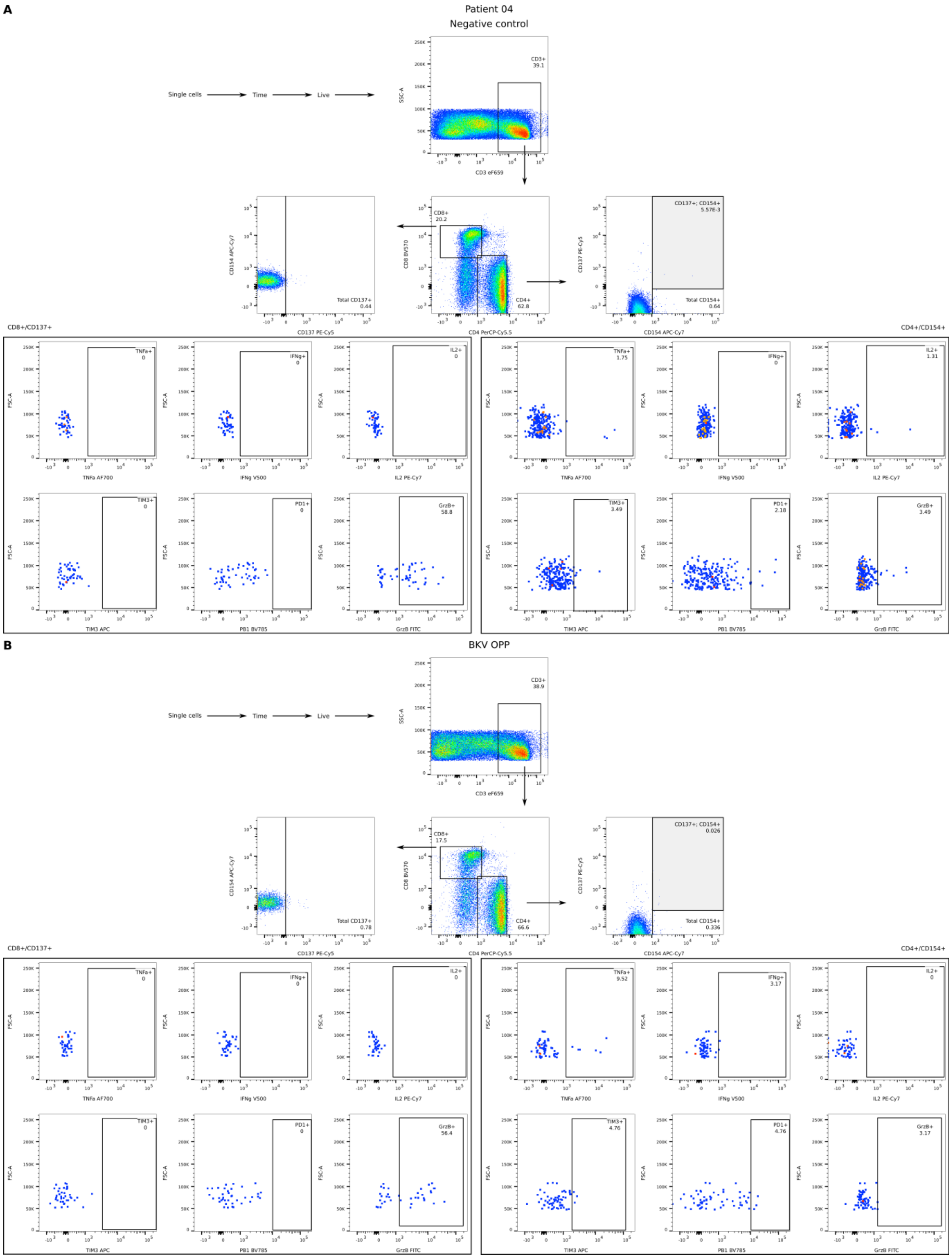


**Supplementary Figure 5.** Gating strategy for the phenotypic and functional characterization of BKV-specific T cells. Complete gating strategy from identification of $CD3^+$ T cells to the assessment of cytokines and exhaustion markers is presented for patient 04. A) Negative DMSO control. B) Stimulation with BKV OPP. For $CD4^+$ $CD154^+CD137^+$ (gray box) and total $CD154^+$ (gray and white box) were identified.

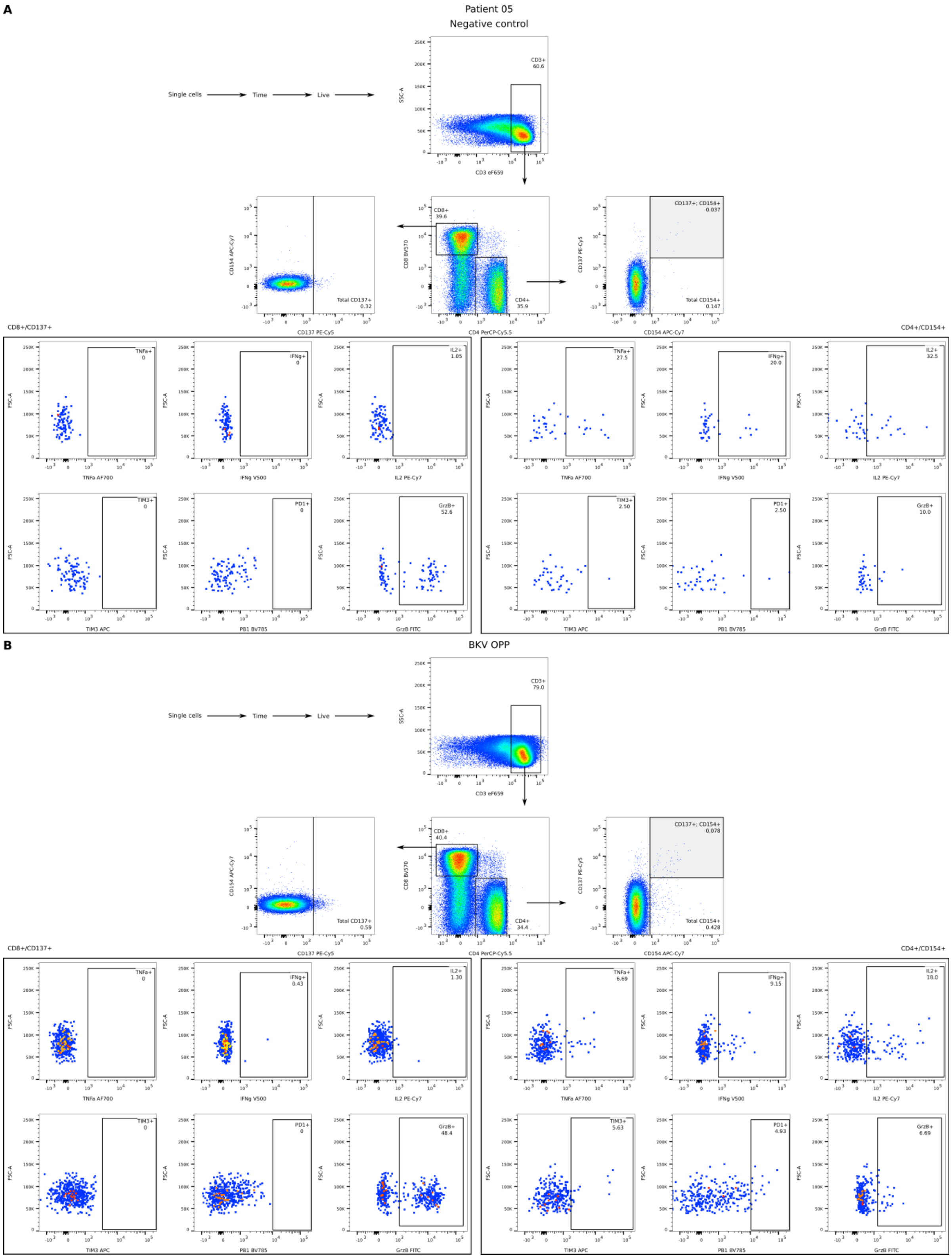


**Supplementary Figure 6.** Gating strategy for the phenotypic and functional characterization of BKV-specific T cells. Complete gating strategy from identification of $CD3^{+}$ T cells to the assessment of cytokines and exhaustion markers is presented for patient 05. A) Negative DMSO control. B) Stimulation with BKV OPP. For $CD4^{+}$ $CD154^{+}CD137^{+}$ (gray box) and total $CD154^{+}$ (gray and white box) were identified.

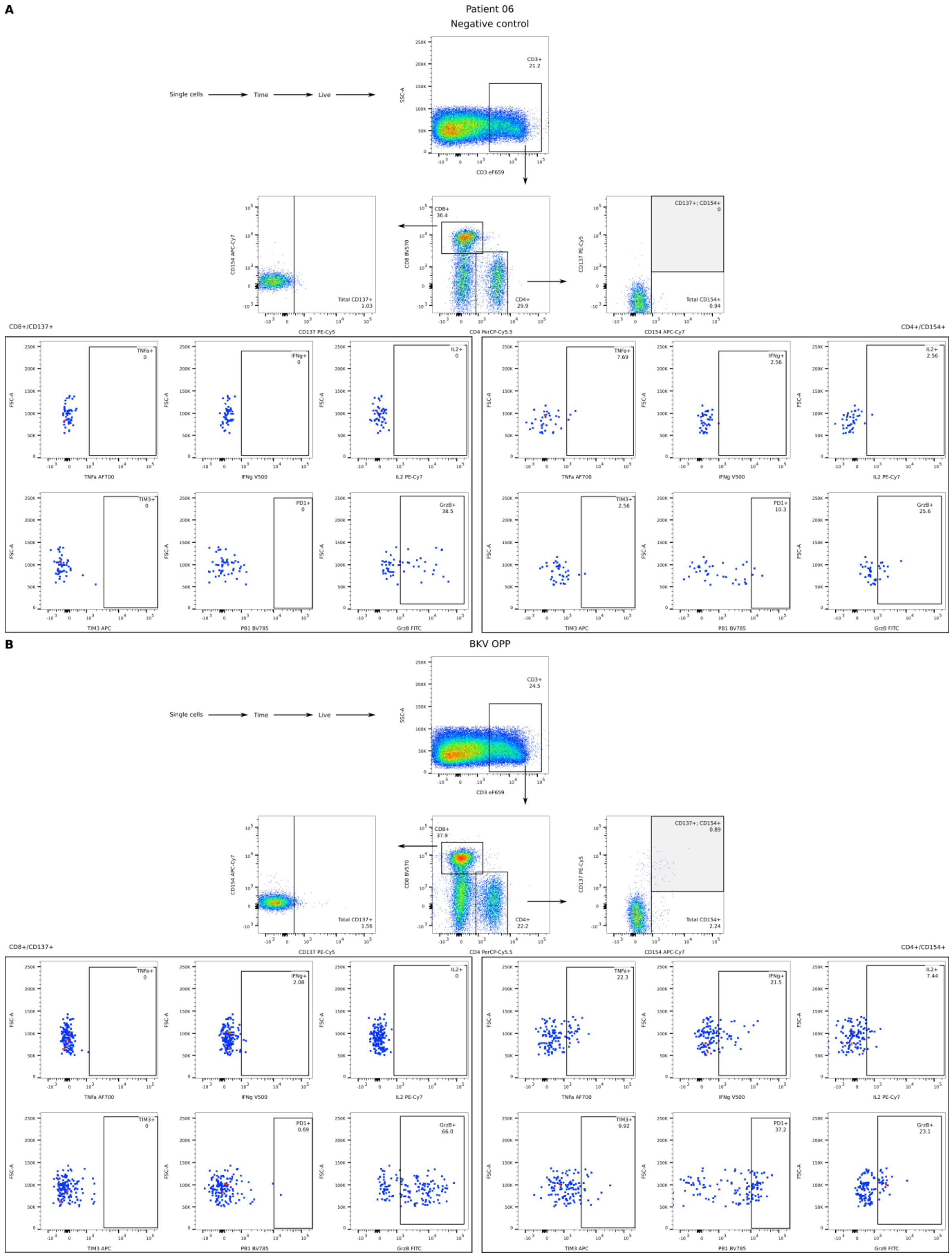


**Supplementary Figure 7.** Gating strategy for the phenotypic and functional characterization of BKV-specific T cells. Complete gating strategy from identification of $CD3^+$ T cells to the assessment of cytokines and exhaustion markers is presented for patient 06. A) Negative DMSO control. B) Stimulation with BKV OPP. For $CD4^+$ $CD154^+CD137^+$ (gray box) and total $CD154^+$ (gray and white box) were identified.

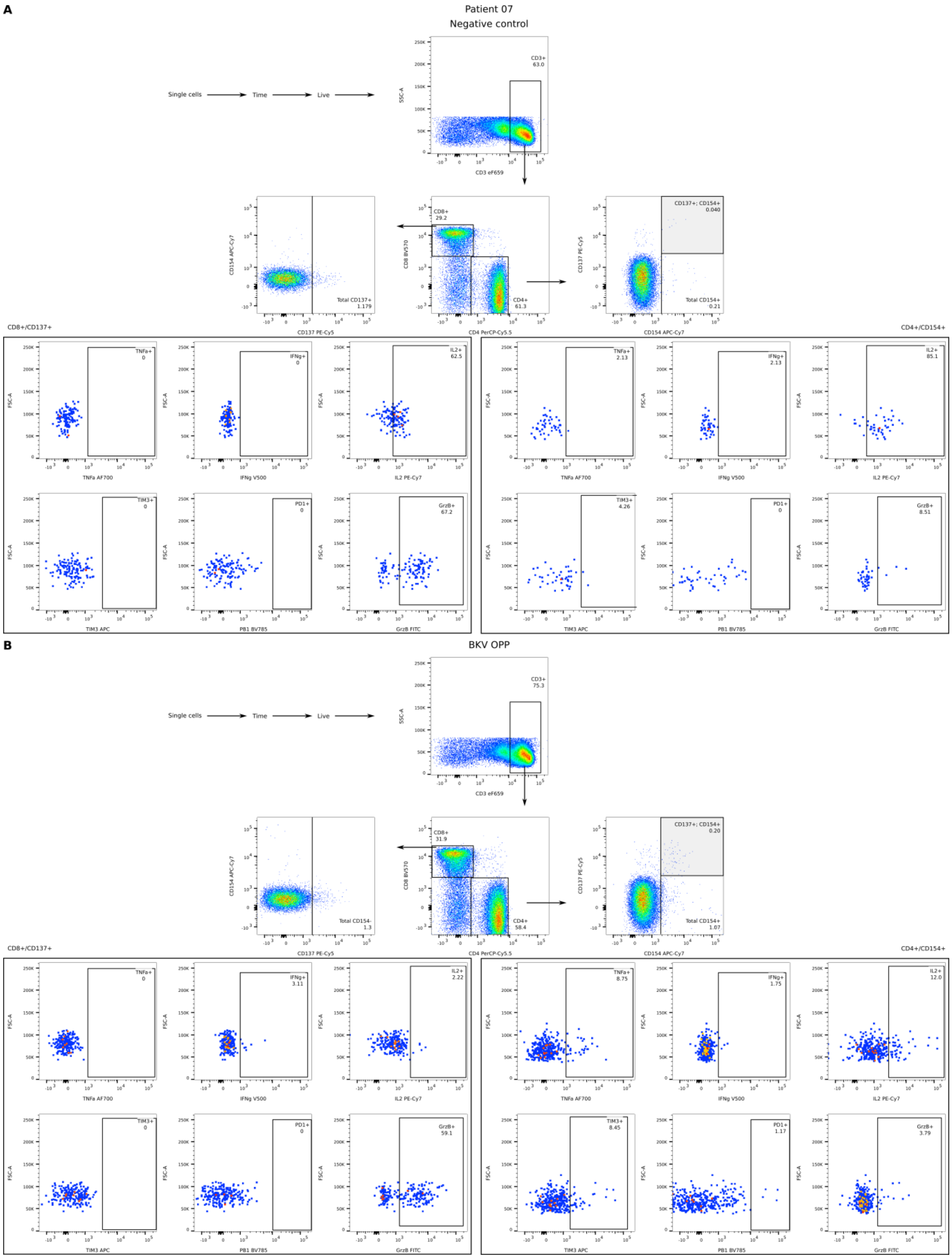


**Supplementary Figure 8.** Gating strategy for the phenotypic and functional characterization of BKV-specific T cells. Complete gating strategy from identification of $CD3^{+}$ T cells to the assessment of cytokines and exhaustion markers is presented for patient 07. A) Negative DMSO control. B) Stimulation with BKV OPP. For $CD4^{+}$ $CD154^{+}CD137^{+}$ (gray box) and total $CD154^{+}$ (gray and white box) were identified.

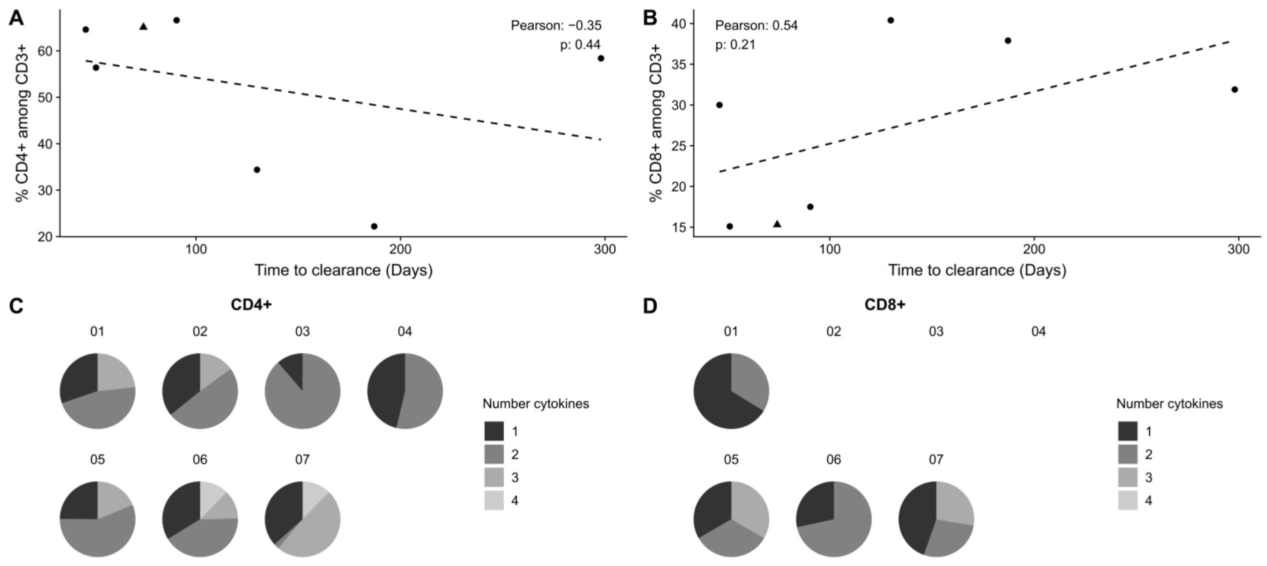


**Supplementary Figure 9.** Correlation between total CD4$^{+}$ (A) and total CD8$^{+}$ (B) T cells and clearance time. Each point represents a patient in the study and the dotted line the best linear fit. C-D) Poly and multi-functionality of CD4$^{+}$/CD154$^{+}$ (C) and CD8$^{+}$/CD137$^{+}$ (D) T cells. For patient 2-4 no cytokines were identified among the CD8$^{+}$/CD137$^{+}$.

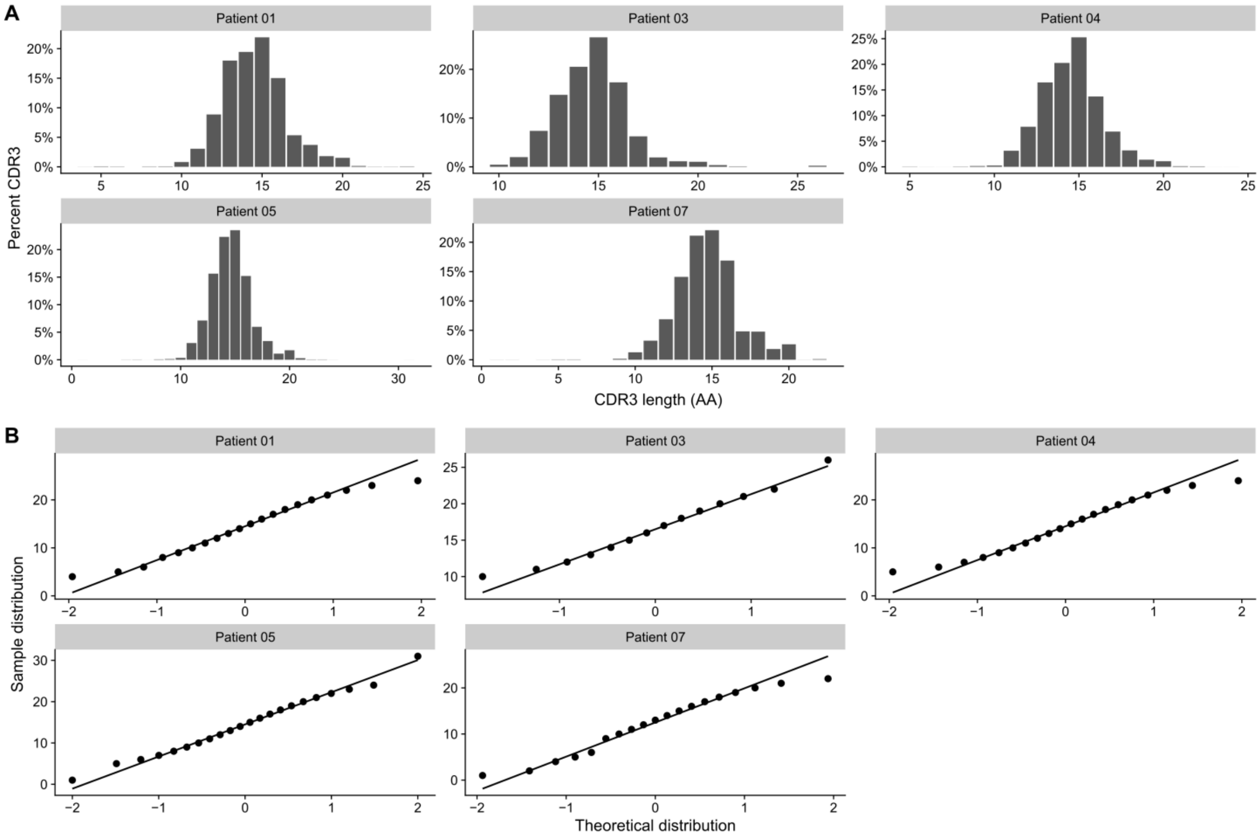


**Supplementary Figure 10.** A) Distribution of TCRβ CDR3 lengths in number of amino acids of BKV-specific TCR clonotypes. The abundance of each length is given as the frequency among all lengths for the patient. B) Quantile-quantile plot comparing the CDR3 length distribution to a standard normal population. The straight line is the best linear fit to the data within the 25th and 75th quantile.

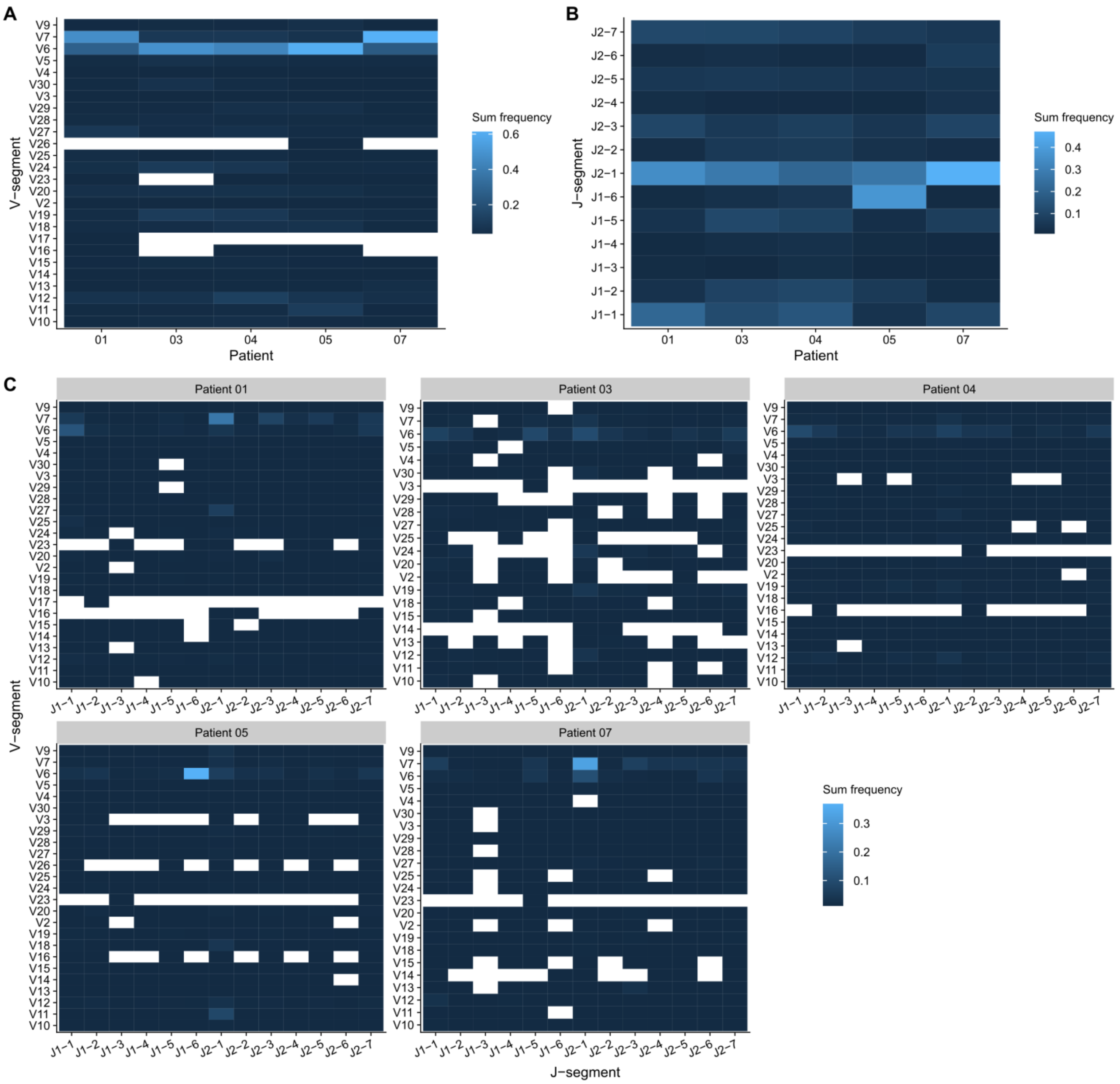


**Supplementary Figure 11.** Distribution of V- (A) and J-segments (B), and their respective combination (C). in BKV-specific clonotypes. The color indicates the total sum of frequencies of TCR clonotypes with a given segment of combination.

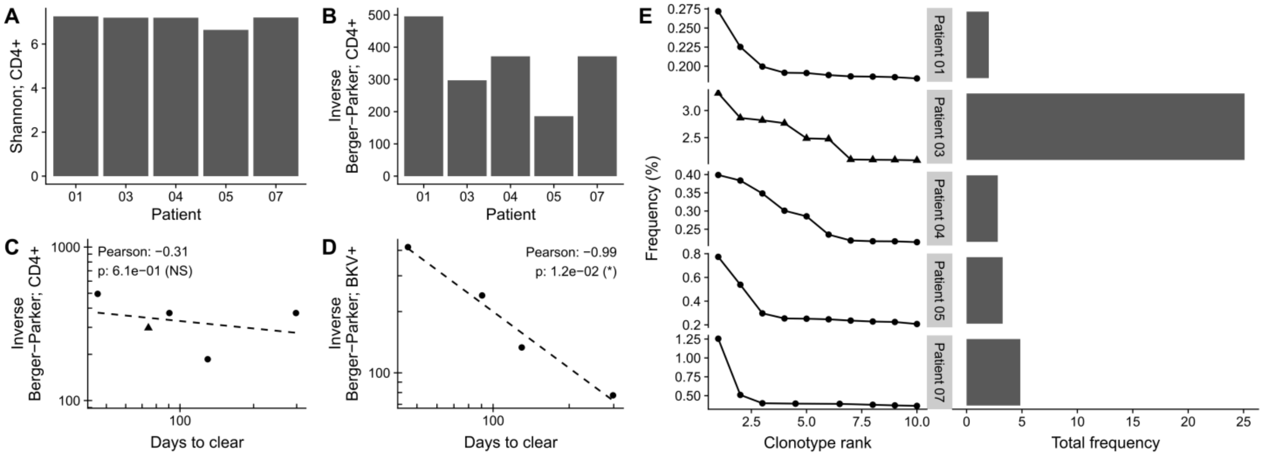


**Supplementary Figure 12.** A) Shannon index for total CD4$^+$ T cells. B) Inverse Berger-Parker for total CD4$^+$ T cells. C) Correlation between inverse Berger-Parker for total CD4$^+$ and clearance time. D) Correlation between inverse Berger-Parker for BKV specific CD4$^+$ and clearance time. E) Rank-abundance of the 10 most dominant clonotypes for each patient (left), and summed abundance (right). Each point in C and D represents a patient in the study and the dotted line the best linear fit.

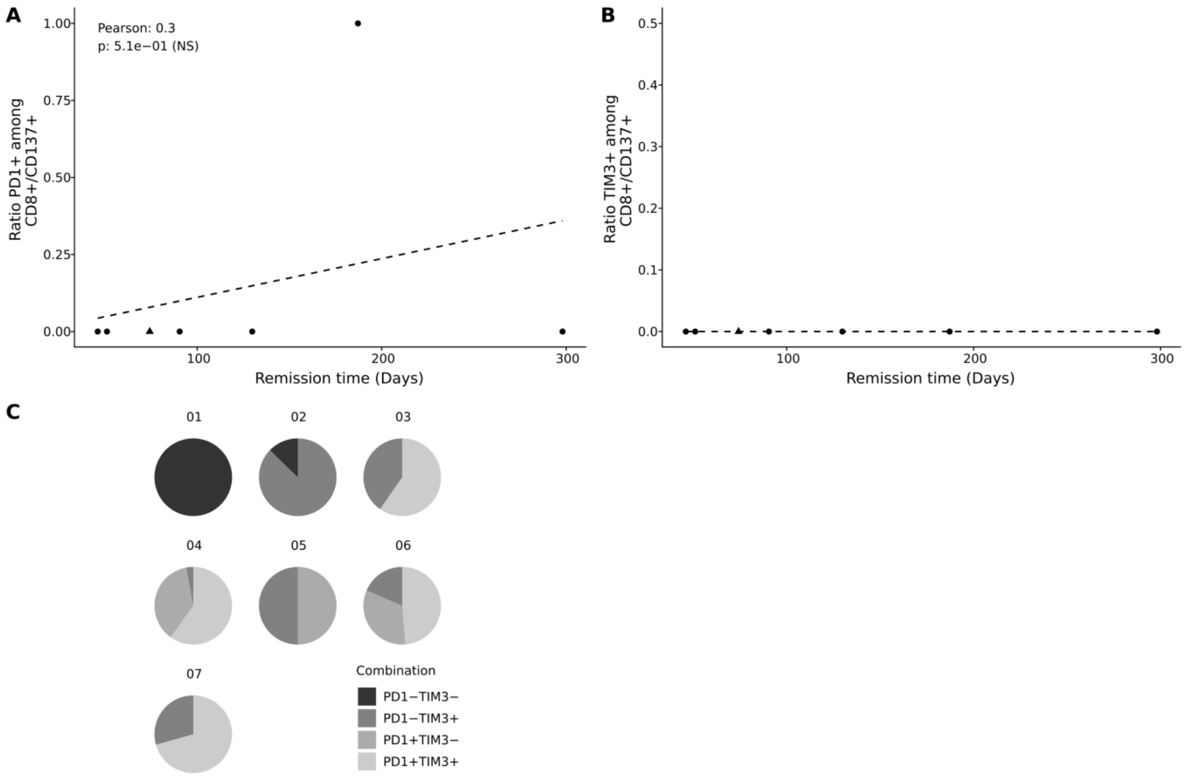


**Supplementary Figure 13.** A-B) Exhaustion markers PD1 (A) and TIM3 (B) on activated $CD8^+$ T cells. Each point indicate a patient, the triangle indicate patient 03 C) Co-expression of PD1 and TIM3 by activated CD4+/CD154+ T cells. Each patient is plotted separately, indicated by the number above the pic chart.

### 1.2 Supplementary Tables

**Supplementary Table 1. Summary statistics of TCRβ CDR3 lengths in number of amino acids.**

| | Length (AA) | | | |
|---|---|---|---|---|
| Patient | Minimum | Average | Maximum | Most likely |
| 1 | 6 | 15 | 24 | 15 |
| 3 | 10 | 17 | 26 | 15 |
| 4 | 6 | 15 | 24 | 15 |
| 5 | 5 | 15 | 31 | 15 |
| 7 | 5 | 15 | 22 | 14 |
